# *Pseudomonas aeruginosa* strains from both clinical and environmental origins readily adopt a stable small colony variant (SCV) phenotype resulting from single mutations in c-di-GMP pathways

**DOI:** 10.1101/2022.06.02.494627

**Authors:** Alison Besse, Marie-Christine Groleau, Mylène Trottier, Antony T. Vincent, Eric Déziel

**Affiliations:** Centre Armand-Frappier Santé Biotechnologie, Institut National de la Recherche Scientifique (INRS), Laval, Québec, H7V 1B7, Canada; Département des Sciences Animales, Faculté des Sciences de l’Agriculture et de l’Alimentation, Université Laval, Québec City, QC, G1V 0A6, Canada; Institut de biologie intégrative et des systèmes, Université Laval, Québec City, QC, G1V 0A6, Canada

## Abstract

A subpopulation of Small Colony Variants (SCVs) is a frequently observed feature of *Pseudomonas aeruginosa* isolated from colonized cystic fibrosis lungs. Since most SCVs have until now been isolated from clinical samples, it remains unclear how widespread is the ability of *P. aeruginosa* to develop this phenotype and the genetic mechanism(s) behind SCVs emergence according to the origin of the isolate. In the present work, we investigated the ability of 22 *P. aeruginosa* isolates from various environmental origins to, under laboratory culture conditions, spontaneously adopt a SCV-like smaller alternative morphotype distinguishable from the ancestral parent strain. We found that all the *P. aeruginosa* strains tested could adopt a SCV phenotype, regardless of their origin. Whole genome sequencing of SCVs obtained from clinical and environmental sources revealed single mutations exclusively in two distinct c-di-GMP signaling pathways, Wsp and YfiBNR. We conclude that the ability to switch to a SCV phenotype is a conserved feature of *P. aeruginosa* and results from the acquisition of a stable genetic mutation, regardless of the origin of the strain.

**IMPORTANCE:** *P. aeruginosa* is an opportunistic pathogen that thrives in many environments. It poses a significant health concern, notably because this bacterium is the most prevalent pathogen found in the lungs of people with cystic fibrosis. In infected hosts, its persistence is considered related to the emergence of an alternative small colony variant (SCV) phenotype. By reporting the distribution of *P. aeruginosa* SCVs in various non-clinical environments and the involvement of c-di-GMP in SCV emergence from both clinical and environmental strains, this work contributes to understanding a conserved adaptation mechanism used by *P. aeruginosa* to adapt readily in all environments. Hindering this adaptation strategy could help control *P. aeruginosa* persistent infection.

## INTRODUCTION

The high genomic and metabolic diversity of *Pseudomonas aeruginosa* allows this bacterium to thrive in diverse environments, such as aquatic habitats, soil, food, and even built environments, such as hospital premise plumbing systems (1-3). This opportunistic pathogen, frequently identified as a causative agent of nosocomial infections, is a major cause of infections in immunocompromised individuals. Notably, *P. aeruginosa* is the most prevalent pathogen found in the lungs of people with cystic fibrosis (CF) (4-6).

*P. aeruginosa* expresses a broad range of virulence determinants that counteract the host immunity and promote survival (7). One of these factors is the ability to form biofilms. These organized communities largely contribute to evade host immunity and antimicrobial treatments. For instance, the biofilm matrix delays penetration of antibiotics and host defense effectors (8-10). *P. aeruginosa* typically persists in the lungs of CF individuals as a biofilm (11, 12).

The emergence of a subpopulation of Small Colony Variants (SCVs) is a frequently observed feature of *P. aeruginosa* isolates from CF lungs biofilms (13, 14). SCVs are characterized by circular opaque dwarf colonies with a diameter about three-time smaller than wild-type (WT) colonies (14-17). Shortly after their first report, we proposed that SCVs are phenotypic variants (18). Several studies suggest that phenotypic switching could be regulated by a reversible adaptation mechanism: phase variation (18, 19), traditionally defined as a high-frequency ON/OFF switch between phenotypes in a heritable and reversible manner (20-22). Indeed, SCVs spontaneously revert to the WT-like morphotype (15, 16, 18, 23). Yet, recent studies have reported stable genetic mutations in *P. aeruginosa* leading to SCV phenotype in *in vitro* grown biofilms and animal model of PA14 infection (14, 24, 25). The SCV phenotype is typically caused by mutations in genes involved in the metabolism of the intracellular second messenger c-di-GMP (14, 26). Among them, mutations in the Wsp (Wrinkly Spreader) pathway are the most frequently reported (14, 24, 27). The Wsp pathway is a chemosensory system resulting in activation of the diguanylate cyclase (DGC) WspR in response to surface sensing, which regulates the c-di-GMP pool, along with other DGCs (synthesis of c-di-GMP) and phosphodiesterases (PDE, degradation of c-di-GMP) in *P. aeruginosa* (28-30).

C-di-GMP is largely involved in regulation of the phenotypic properties associated with SCVs, though binding to specific receptors. For instance, while an overproduction of exopolysaccharides (EPS) (Pel and Psl) (14, 31) and a motility deficiency, notably flagellar, has been described for SCVs (16, 18, 32), high c-di-GMP levels activate the expression of the *pel* operon, leading to production of the EPS Pel, and repress flagellar motility (33-35). *P. aeruginosa* SCVs exhibit several other specific properties such as cell surface hyperpiliation and adherence to abiotic surfaces (16, 18, 36). These properties promote biofilm formation (37). Additionally, SCVs exhibit autoaggregative properties (16, 36).

It is striking that SCVs have been mostly isolated from infected hosts, essentially CF individuals; or by extension, from laboratory cultivation of strains sampled from infected hosts (13). For instance, several studies have recovered SCVs from lung, sputum or deep throat swabs of CF individuals (12, 16, 17, 38). CF is not the only pathology associated with the emergence of *P. aeruginosa* SCVs. These variants have also been isolated from urine, feces, endotracheal secretion and pleural effusion of patients suffering from meningioma, anoxic encephalopathy, hepatocellular carcinoma, lung carcinoma or grave asphyxia neonatorum (39). In addition to having been isolated from infected hosts, SCVs have also been generated under *in vivo* laboratory conditions. For instance, SCVs have been obtained *in vivo* from *P. aeruginosa* strains during infections in burn wound porcine models and murine models (24, 40). In the latter study, the authors have clearly showed that SCVs arisen in infection context are due stable genetic mutation in their genomes (24).

Intriguingly, 20 years ago we reported one of the first identification of *P. aeruginosa* SCVs that quickly emerged when a soil isolate was grown on a non-aqueous phase liquid, hexadecane, as sole substrate (18). The SCV morphotype of strain 57RP predominates when biofilm growth conditions are preferable and displays features shared with clinical SCVs: high adherence, efficient biofilm formation, hyperpiliation and reduced motility (18).To our knowledge, this study is the only one reporting SCVs for an environmental *P. aeruginosa* isolate. However, the genetic cause leading to SCV emergence in the environmental context remains elusive. SCVs generated *in vitro* from *P. aeruginosa* PAO1 and PA14 showed stable mutations, but these strains, although prototypical, are still of clinical origin (25, 36).

Since most SCVs have until now been isolated from clinical samples, it remains unclear how widespread is the ability of *P. aeruginosa* to develop this phenotype and the genetic mechanism(s) behind SCVs emergence in regard to the origin of the isolate: are they exploiting phase variation or selecting adaptive mutants? Here, we investigated the ability of *P. aeruginosa* isolates from various environmental origins to spontaneously adopt, under laboratory culture conditions, a SCV-like smaller colony morphotype readily distinguishable from their ancestral parent. We tested 22 *P. aeruginosa* strains from four different categories of environments: soil, food, hospital water systems and clinical; we found that all the *P. aeruginosa* strains have the ability to adopt the SCV phenotype, regardless of their origin. Whole genome sequencing was performed on SCVs from two strains isolated from distinct environments to investigate the potential genetic causes responsible for the SCV phenotypes. We found that mutations affecting c-di-GMP signalling pathways were responsible for SCV emergence in clinical and environmental strains.

## RESULTS

### The ability to form SCV-like morphotype colonies is a conserved feature of *Pseudomonas aeruginosa*

Culture conditions promoting biofilm formation select for SCVs of *P. aeruginosa* (16, 18, 36). To broadly investigate the ability of *P. aeruginosa* to adopt a SCV-like morphotype, we cultured 22 isolates from various origins in static liquid medium for 65 h then spread onto TSA plates to obtain isolated colonies. Six strains were from food samples (meat and fish from markets), six from clinical samples (five from CF patients and the clinical prototypic strain PA14 from a burn patient), five from petroleum oil-contaminated soil and five from hospital sinks (drain, splash area and tap) (Table 1, columns 1 and 2). To cover the variety of temperatures relevant to these various habitats, the cultures were incubated in a temperature range varying from 30 to 40°C. At the onset, none of the strains were displaying a SCV phenotype, but after 65 h of incubation all isolates diversified in a range of colony morphotypes, including small colonies that appeared typical of SCVs (Fig. 1, for selected strains from each origin). Small colonies emerged in the cultures incubated at all tested temperatures (data not shown).

**Table 1.**
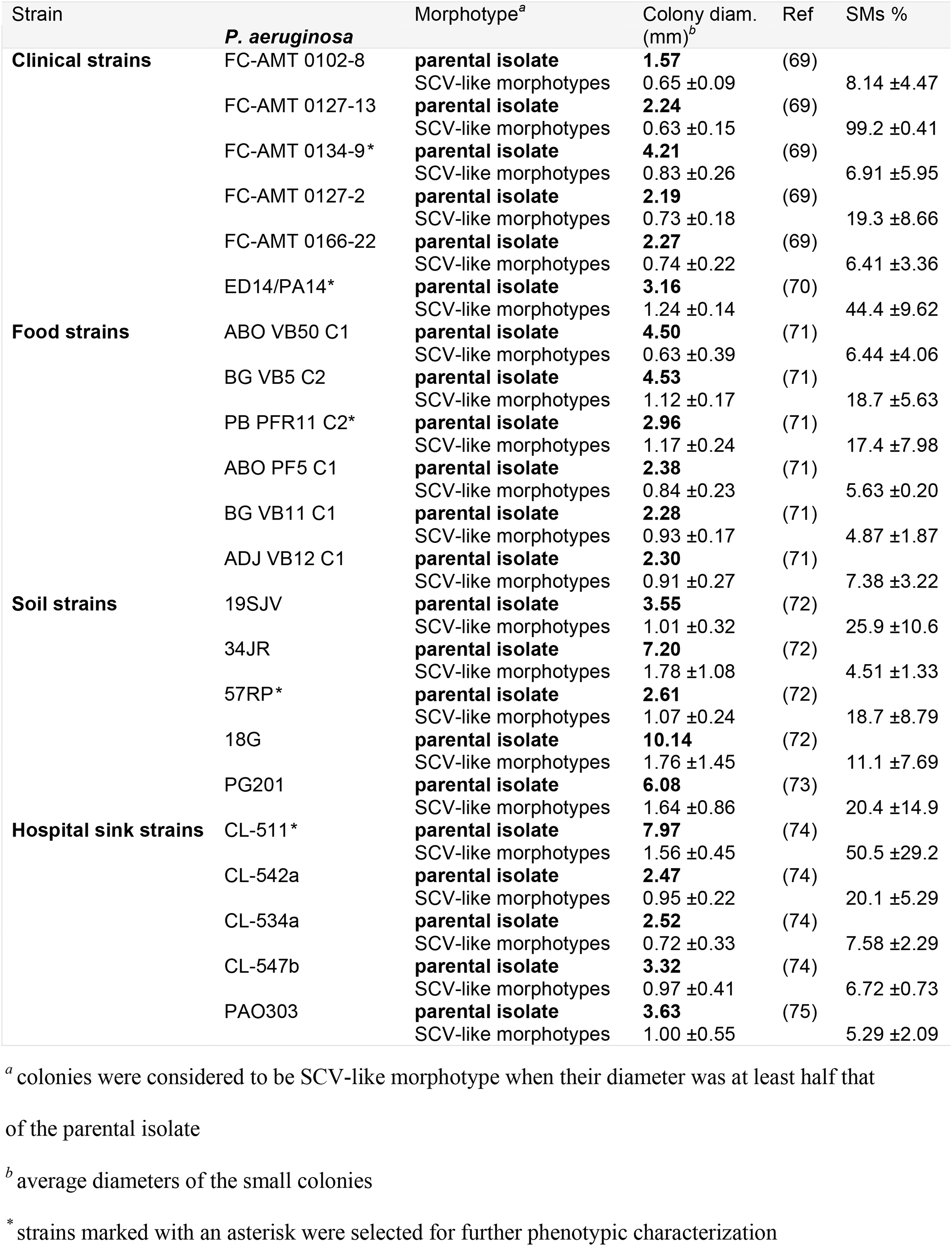
Colony diameters and prevalence of parental isolates and their static liquid culture evolved small morphotypes.

**Fig. 1.**
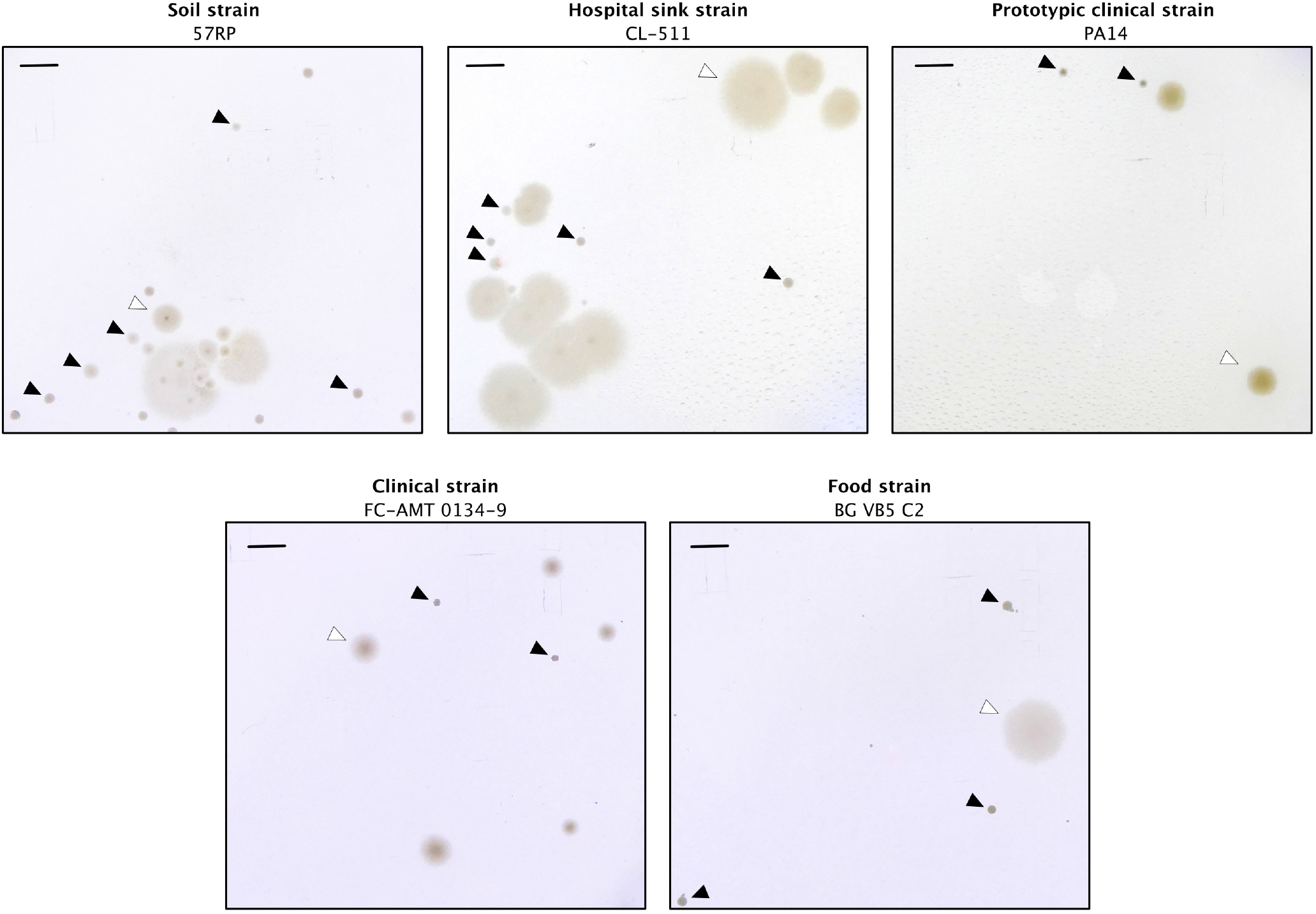
Small colonies of *Pseudomonas aeruginosa* emerge in static cultures from strains isolated from various origins. Parental strains were inoculated under static liquid conditions in TSB for 65 hours and spread onto TS-Agar 2% plates. Black arrows indicate smaller colonies. White arrows indicate parent-like colony. Scale bars represent 5 mm.

Reported SCVs have an average diameter two to four times smaller than WT colonies. Colonies correspondingly smaller than the parental strains emerged from all 22 strains (Table 1). This result strongly suggests that the ability to produce variant colonies displaying an SCV-like morphotype is a conserved feature of *P. aeruginosa*, regardless of the origin of the strains.

### Isolated SCV-like morphotype colonies belong to two distinct clusters

By taking a closer look at the emerged SCV-like morphotypes, we observed that their sizes (Table 1) and overall appearance (Fig. 1) differ. Some colonies were denser, with well-defined round edges and others were more translucent with undefined edges (Fig. 1). We then asked whether these different types of SCV-like morphotypes are indeed *bona fide* SCVs, and if a distinction can be made between them. We focused on five strains from different origins, (Table 1, strains indicated by an asterisk) and isolated the various distinct morphotypic small colonies (SMs for SCV-like morphotype) produced by each following static incubation and plating. Besides their sizes, we looked at several phenotypes typically associated with SCVs: swimming motility defect, biofilm formation and production of EPS, cell aggregation and production of c-di-GMP. Because cell aggregation induces the production of pyoverdine, the fluorescent siderophore of *P. aeruginosa*, while loss of the EPS coding genes, *pel* and *psl*, leads to inhibition of pyoverdine production (41), we used the production of pyoverdine as an indirect measurement of cell aggregation and EPS production. We compiled the phenotypical data for each distinct SMs (Table S1) and performed a principal coordinates analysis (PCoA) based on their colony size, auto-aggregation properties (pyoverdine production), their ability to perform swimming motility, timing of biofilm formation and total biomass of biofilms. In a PCoA, all variables are equally considered to cluster SCVs in significant groups based on their phenotypic profiles and to better understand which SMs are close to each other and could be part of the same clusters. We found that the various distinct SMs generated by the five parental strains clustered in two separate groups (named Cluster 1 and Cluster 2) (Fig. 2). Members of both clusters for the SMs of soil strain 57RP, the sink hospital strain CL-511, the food strain PB PFR11 C2, and the clinical strain FC-AMT0134-9 had phenotypical features that distinguished them from their parental strain (Fig. 2). Cluster 2 of strain PA14 contained only one isolated SM, but we believe that this is only the result of lower abundance of this form when sampling was performed. These results indicate that two distinct phenotypic types of SCV-like morphotypes emerged under our culture conditions.

**Fig. 2.**
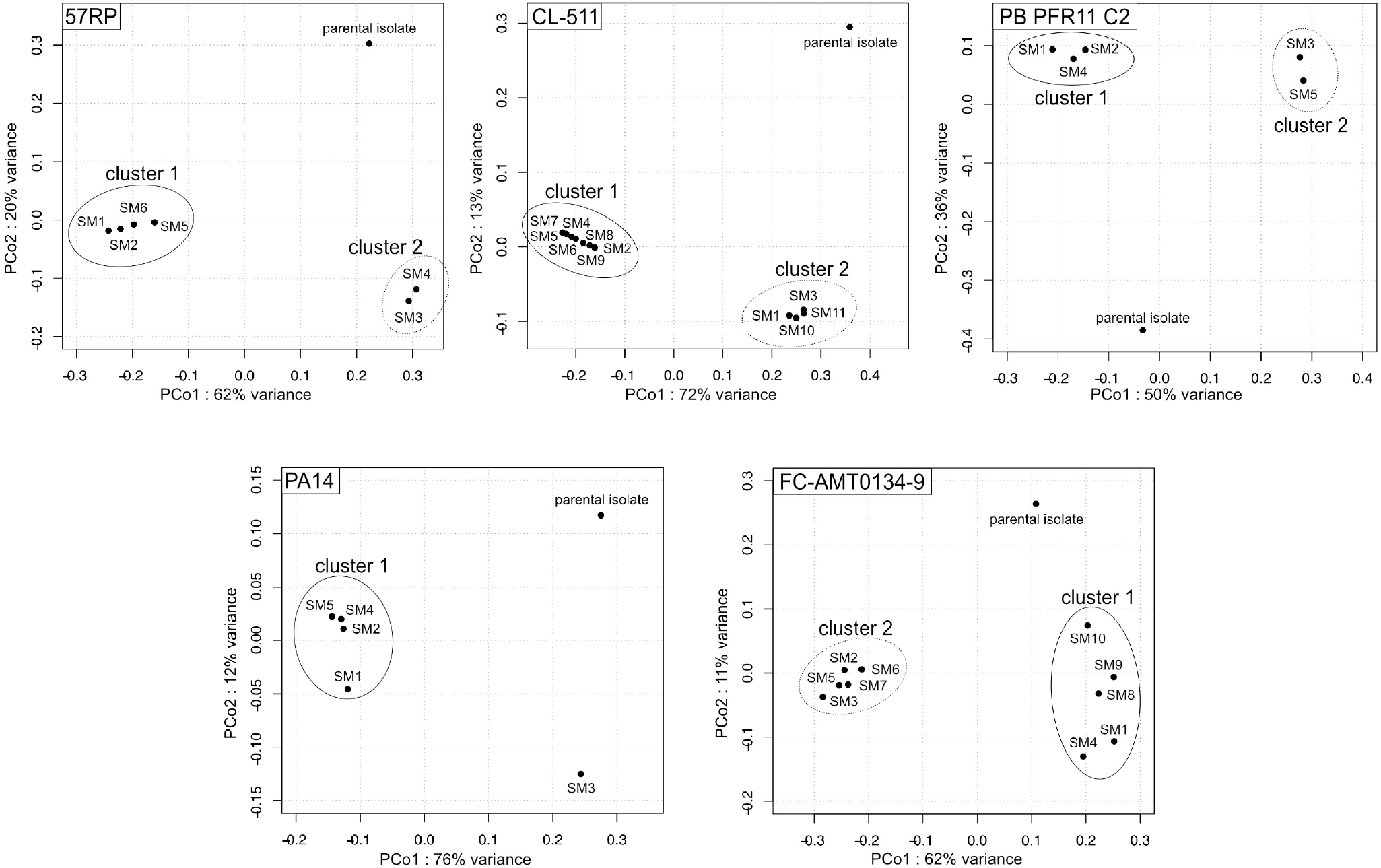
Small colonies isolated from static cultures are clustered in two separate groups according to their phenotypic features. PCoA analysis were performed with a matrix composed of data obtained from the phenotypic analyses (swimming, biofilm formation, and pyoverdine production) for the parental strain and distinct small colonies isolated from static cultures with a diameter at least two times smaller than parental strain (Table S1). All variables were equally considered to cluster colonies in significant groups based on their phenotypic profiles. Each point represents a small colony isolated from the static cultures and has a name code composed of SMx standing for Small Morphotype where x is an arbitrary number attributed during the isolation of the colonies. The identification of statistically distinctive clusters was performed using simprof tests and hclust.

### SMs from Cluster 1 are typical SCVs with a reversible state

SMs belonging to Cluster 1 of each strain share some common features: a reduced swimming motility, and/or a promoted biofilm formation, and/or enhanced auto-aggregation properties (pyoverdine production) as compared with their parental strain (Table S1 and Fig. S1). These features are typical of SCVs described in the literature. Since these phenotypes are regulated by c-di-GMP, we assessed intracellular c-di-GMP levels in selected SMs of Cluster 1. As expected, higher c-di-GMP levels were measured in Cluster 1 SMs than in their parental counterparts, again indicating that Cluster 1 SMs are indeed typical SCVs (Fig. 3). In addition to quantitative PCoA data, we looked at rugosity of SM colonies, a qualitative phenotype traditionally associated with SCVs. While Cluster 1 SMs colonies display a very distinctive rugose surface as compared with their parental counterparts, rugosity appearance was diverse among the strains (Fig. 4). In conclusion, phenotypic characterisation confirms that SMs belonging to Cluster 1 are typical SCVs.

**Fig. 3.**
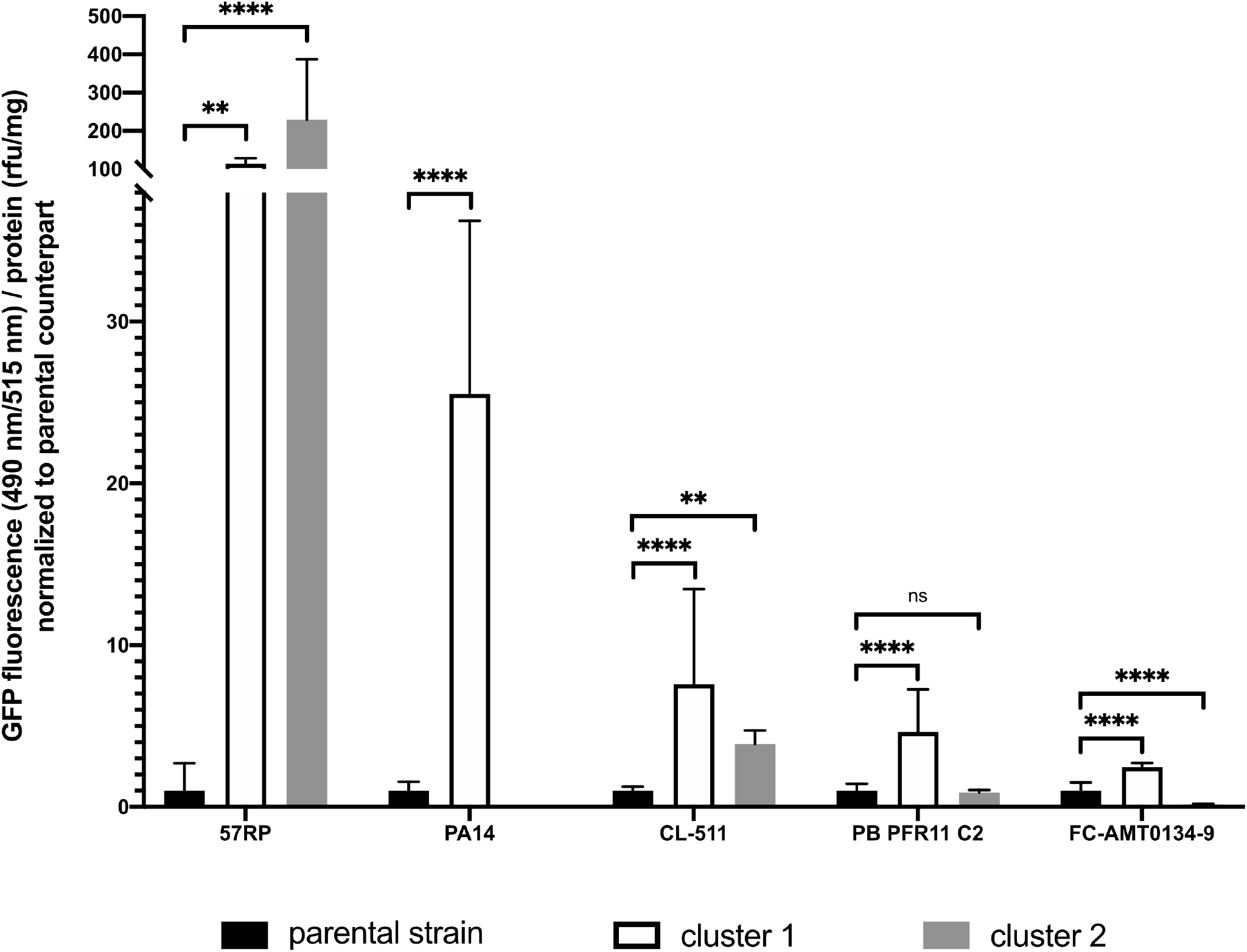
c-di-GMP production is altered for SMs from Cluster 1 and 2 compared with their respective parental strain. c-di-GMP production was measured with the fluorescent-based biosensor pCdrA-gfp on O/N washed cultures. The values are means ± standard deviations (error bars) for selected transformed morphotypes belonging to each cluster: transformed morphotypes were SM2 and SM6 (cluster 1) and SM4 (cluster 2) for strain 57RP; SM4 and SM5 (cluster 1) for strain PA14; SM8 and SM9 (cluster 1) and SM10 (cluster 2) for strain CL-511; SM1 and SM2 (cluster 1) and SM3 and SM6 (cluster 2) for strain PB PFRC11 2; SM9 (cluster 1) and SM5 and SM7 (cluster 2) for strain FC-AMT0134-9. Three transformants of each SMs were considered in the calculation of mean and standard deviations. Stars represents the statistical significance of the results calculated by an Ordinary one-way analysis of variance (ANOVA), ****, P Value ≤ 0.0001; **, P Value ≤ 0.01; ns, not significant. Data are normalized between them based on their parental strain.

**Fig. 4.**
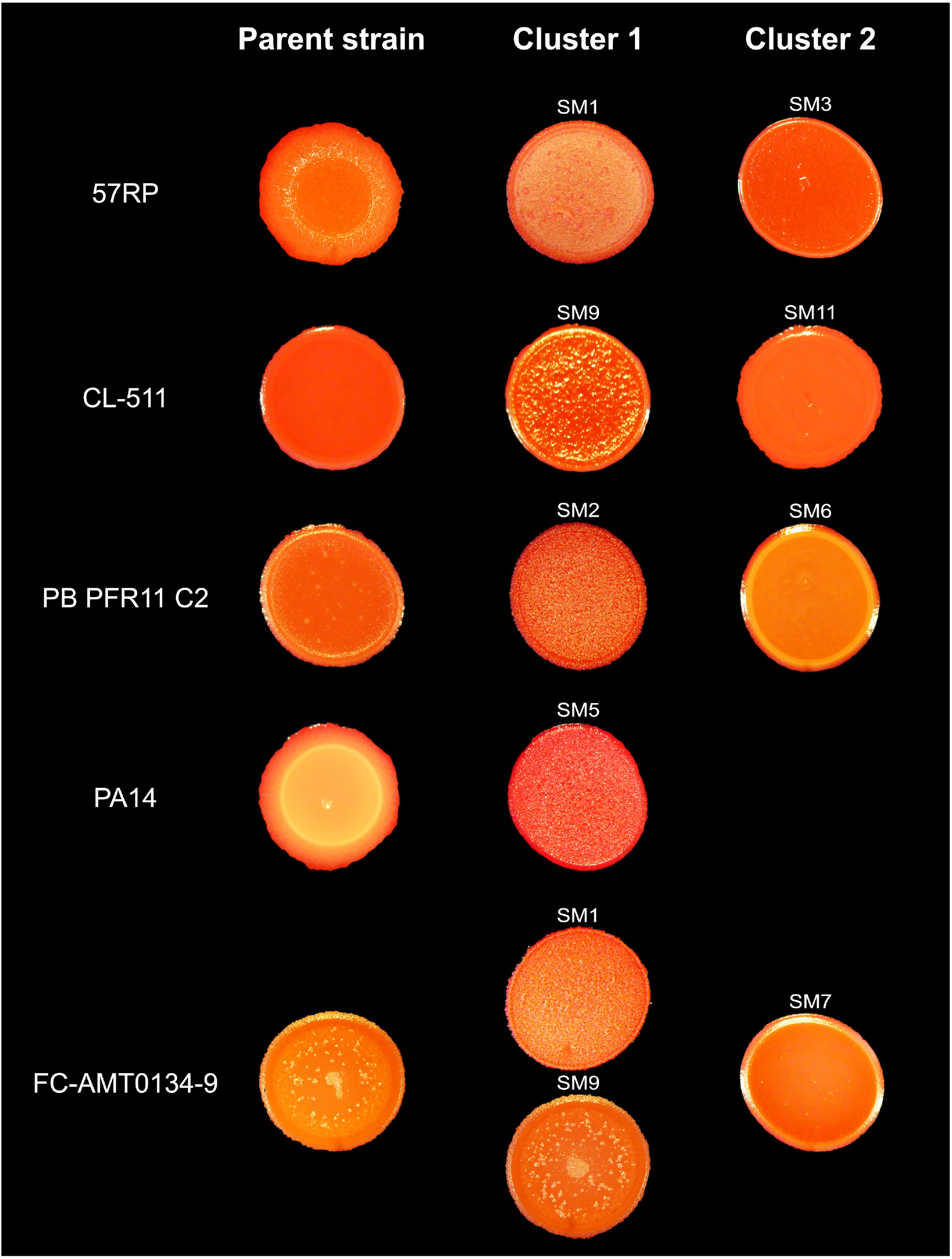
Appearance of colonies for the parental isolates and SMs from Cluster 1 and Cluster 2 on Congo Red plates. The SM shown for each cluster is representative of all the SMs included in one cluster since they have a similar appearance. Plates were observed with a binocular StemiDV4 (Zeiss) and photos were taken with a DMC-ZS60 camera (Panasonic Lumix), after 24 h of incubation at 30°C.

Finally, we observed the emergence of spontaneous reversion to a larger, parental-like phenotype, a property typically associated with phase variation. As stated above, on agar plates, reversion to the parental-like morphotype was observed after a 48h incubation at 30°C for SMs belonging to Cluster 1 (Fig. 5). Reversion was revealed as an outgrowth from the original colony, but sometimes only by a change in appearance of the colony surface, as seen for instance with isolate PB PFR11 C2 (Fig. 5). This reversibility suggested that SCVs could arise from a phase variation process.

**Fig. 5.**
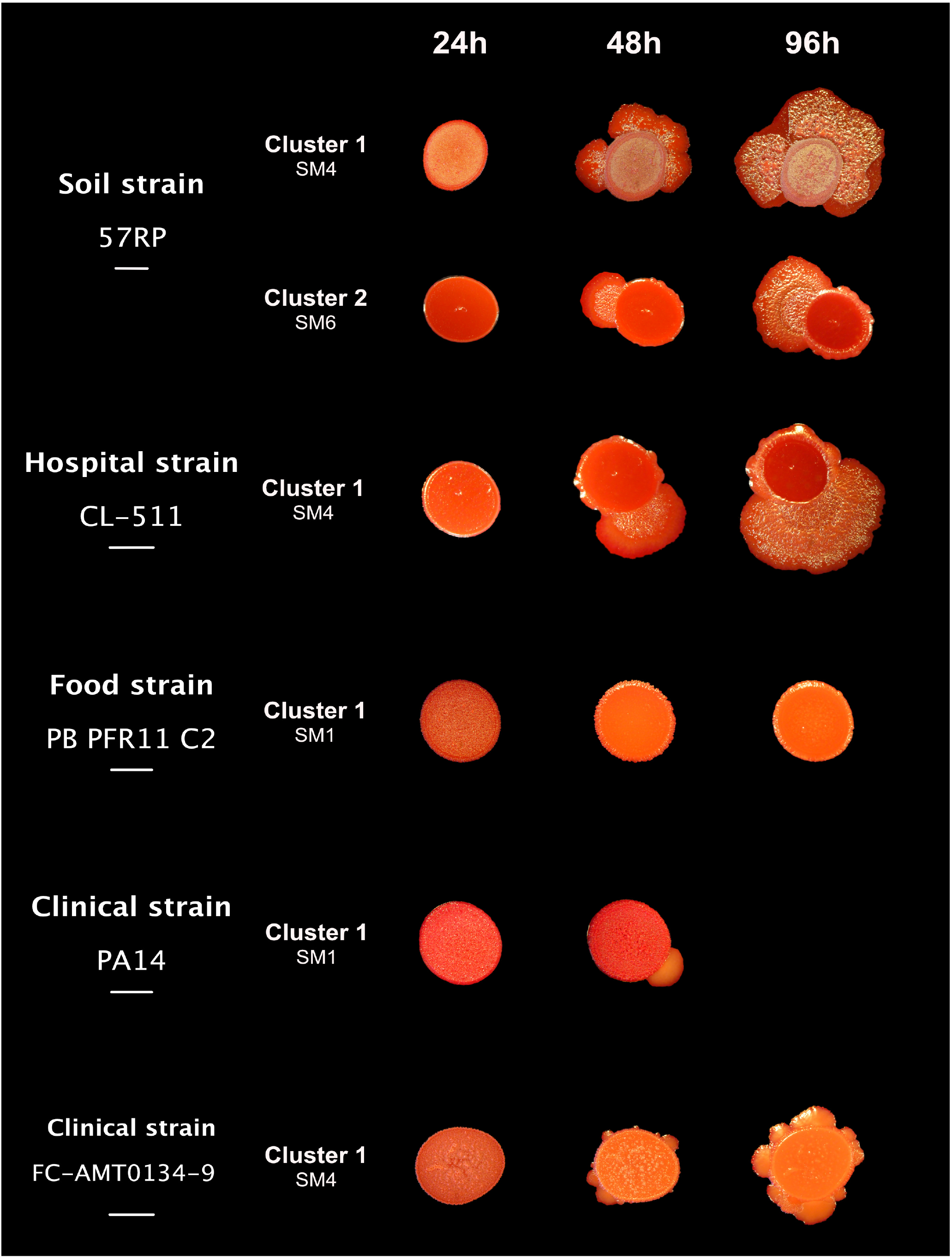
Reversion occurs on solid media for specific morphotypes after 48 h incubation. Ten µl of a culture of parental strain or a cluster representative morphotype (SMs) were dropped on 0.1% congo red TS-Agar 2% plates. Plates were observed with a binocular StemiDV4 (Zeiss) and photos were taken with the camera DMC-ZS60 (Panasonic Lumix), after 24 h, 48 h and 96 h of incubation at 30°C. Scale bars represent 5 mm.

### Cluster 1 SCVs harbor mutation in c-di-GMP pathways

To investigate if SCVs could arise from phase variation, we performed whole genome sequencing of PA14 and 57RP SCVs obtained from independent experiments of 65h static cultivation with the parental strains. The genomes of the parental strains were used as reference for the search for potential mutations in SCVs genomes. Mutations were found in all SCVs (Table 2). Interestingly, they were exclusively detected in genes involved in c-di-GMP metabolism. SCVs randomly selected from the first experiment with the parental strain PA14 carry missense single nucleotide polymorphism (SNP) mutations in the *yfiN* gene, while SCVs obtained from the second and third experiment have mutations in the *wsp* cluster, specifically in the *wspA* and *wspF* genes (Table 2). Mutations in the *wsp* cluster are SNPs, resulting in a missense or a stop codon; except for one variant showing a single base deletion leading to a frameshift. SCVs obtained from 57RP carry mutations exclusively in the *wspA* gene; specifically, an in-frame 42-bp deletion (Δ285−298 aa) is present in 12 sequenced SCVs over 13 total (Table 2). The other sequenced 57RP SCV also carry a mutation in *wspA* but it is a missense SNP leading to the replacement of a proline residue by a serine residue, potentially also resulting in modulation of WspA activity (Table 2). These mutations in PA14 and 57RP SCVs genomes are likely to be responsible for the increased c-di-GMP levels we measured (Fig. 3). These results indicate that SCV emergence is largely due to mutations resulting in increased c-di-GMP levels. On the other hand, transposon mutants of *wspR* or *yfiN*, resulting in the inactivation of DGCs WspR or YfiN in PA14 do not affect the rate of SCV emergence, suggesting this phenomenon is regulated by interchangeable DGCs (Fig. S2). Yet, the genetic cause of the SCV phenotype remains elusive: is it always arising from stable mutations or is it a consequence of reversible mutations accounting for the reversion stated above?

**Table 2.** Mutations identified in 57RP and PA14 SCVs and their revertant.

| SCV sequencing<br>(reference genome = parental) |  |  |  |  | Revertant sequencing<br>(reference genome = parental) |  |  |
| --- | --- | --- | --- | --- | --- | --- | --- |
| Strain | Exp. no.<sup>a</sup> | SCV no. | Gene | Mutation<sup>b</sup> | Name | Gene | Mutation |
| **57RP** | 1 | 1 | *wspA* | del 285-298 | **Revertant- SCV**<br>**no. 2** |  |  |
|  |  | 2 | *wspA* | del 285-298 |  | *wspA* | del 285-298 |
|  |  | 3 | *wspA* | del 285-298 |  | *wspR* | Leu71ms |
|  |  | 4 | *wspA* | del 285-298 |  |  |  |
|  |  | 5 | *wspA* | Pro479ms |  |  |  |
|  | 2 | 6 | *wspA* | del 285-298<sup>c</sup> |  |  |  |
|  |  | 7 |  | del 285-298<sup>c</sup> |  |  |  |
|  |  | 8 |  | del 285-298<sup>c</sup> |  |  |  |
|  | 3 | 9 | *wspA* | del 285-298<sup>c</sup> |  |  |  |
|  |  | 11 |  | del 285-298<sup>c</sup> |  |  |  |
|  |  | 12 |  | del 285-298<sup>c</sup> |  |  |  |
|  | 4 | 13 | *wspA* | del 285-298<sup>c</sup> |  |  |  |
|  |  | 14 |  | del 285-298<sup>c</sup> |  |  |  |
|  |  | 15 |  | del 285-298<sup>c</sup> |  |  |  |
| **PA14** | 1 | 1 | *yfiN* | Cys166ms | **Revertant- SCV**<br>**no. 2** |  |  |
|  |  | 2 | *yfiN* | Cys166ms |  | *yfiN* | Cys166ms |
|  |  | 3 | *yfiN* | Cys166ms |  | *yfiN* | Asp304ms |
|  | 2 | 4 | *wspF* | Val318fs |  |  |  |
|  | 3 | 5 | *wspF* | Gln297ns |  |  |  |
|  | 4 | 6 | *wspA* | Ala422ms |  |  |  |

<sup>a</sup> Experiment number – each experiments are performed individually from a distinct inoculum of the parental strain
<sup>b</sup> deleted amino acid residues are indicated. del, deletion; ms, missense; fs, frameshift; ns, nonsense
<sup>c</sup> deletion was detected by PCR and whole genome was not sequenced

### Cluster 1 SCV emergence is due to stable mutations and a second mutation is responsible for reversions

Despite the presence of mutations in the SCVs, reversion is still systematically observed on agar plates; this keeps suggesting that their emergence could be regulated by a phase variation mechanism. For each strain tested, reversions of SCVs are obtained directly within the colony by extending the incubation time of the plate (Fig. 5). However, the inoculation of SCVs under non-favorable conditions, *e*.*g*. cultivation with agitation, did not enable to detect the emergence of revertants, regardless of the strain (data not shown). This result suggests that the mutations that occurred in PA14 and 57RP SCVs were rather stable. To determine if the SCV phenotype was due to a stable or a reversible genetic mutation, whole genome sequencing was performed on reversion outgrowths of SCV PA14 (SM2) and 57RP (SM2) colony (Fig. 5 and Table 2). In the PA14 SCV (SM2) outgrowth, a second SNP mutation was detected downstream of the first mutation in the same gene, *yfiN*, which resulted again in a missense codon. We suppose that this second mutation counterbalances the effect of the first mutation and is responsible for the switch from SCV to another morphotype, probably by inactivating YfiN. In the 57RP SCV (SM2) outgrowth, a second mutation was also detected. However, this mutation is in a different gene, *wspR*, located functionally downstream of the mutated WspA. Thus, regardless of the origin of the strain, reversion was due to a second mutation, indicating that the SCV phenotype is due to the acquisition of a stable genetic mutation and reversion was not the result of phase variation (Fig. 5 and Table 2).

### SMs from Cluster 2 display phenotypical heterogeneity

Unlike Cluster 1 SMs, SMs included in Cluster 2 display inter-strain diversity when considering the phenotypes used for the PCoA (Table S1 and Fig. S1). For instance, among Cluster 2 SMs, swimming motility was intermediate between the parental strain and Cluster 1 SMs for strains 57RP and PB PFR11 C2 (Table S1 and Fig. S1, A). However, for strains CL-511 and FC-AMT0134-9 the swimming motility was increased compared to both Cluster 1 SMs and the parental strains (Table S1 and Fig. S1, A). In addition to PCoA data, c-di-GMP production in Cluster 2 SMs was also variable depending on the parental strain: 57RP Cluster 2 SMs showed higher levels of c-di-GMP compared with both parental strain and Cluster 1 SMs but CL-511 Cluster 2 SMs showed higher production of c-di-GMP only compared to the parental strain (Fig. 3). Also, Cluster 2 SMs in the food strain PB PFR11 C2 showed similar production of c-di-GMP and Cluster 2 SMs in the clinical strain FC-AMT0134-9 even lower production of c-di-GMP as compared to their parental strain (Fig. 3). Thus, c-di-GMP levels are not a consistent driving feature for SMs belonging to Cluster 2. The appearance of the colony surface of Cluster 2 SMs is also distinct on Congo Red plates, once again depending on the parental strain. Colonies of SM3 and SM4 from 57RP display a rugose surface, however less pronounced than for Cluster 1 morphotypes (SM1, SM2, SM5 and SM6), in agreement with their reduced autoaggregative properties (Fig. 4 and Fig. S1, D). For the other strains (PA14, PB PFR11 C2, CL-511 and FC-AMT0134-9), SMs from Cluster 2 display a smoother surface on Congo Red agar, closer to the parental strain (Fig. 4). While Cluster 2 SMs show rapid emergence to reproducible phenotypes, reversion to a larger colonial morphotype akin to WT was only observed for 57RP Cluster 2 SMs and not for the other strains, after 96 h (Fig. 5). All together, these results indicate that, apart from strain 57RP, SMs from Cluster 2 do not exhibit most of the typically described features of SCVs.

## DISCUSSION

### The ability to switch to the SCV phenotype is a conserved feature among *P. aeruginosa* strains, regardless of their origin

SCVs have been mostly reported in the context of human infections, notably from CF individuals. A correlation between the emergence of *P. aeruginosa* SCVs and infection persistence in animal models was established, supporting the idea that the SCV phenotype confers a fitness advantage under chronic infection conditions (42-44). Switch towards the SCV morphotype may represent an adaptation strategy to the hostile environment of the host by increasing resistance to host immunity and antimicrobial treatments (43, 45). However, the emergence of SCVs is not exclusively related to a clinical context. For instance, in 2001 Déziel *et al*. (18) reported the emergence of SCVs in laboratory cultures of a soil *P. aeruginosa* isolate. However, since then, apart from laboratory-grown prototypical strains *P. aeruginosa* PAO1 and PA14, both of clinical origins, no SCVs have been reported from a non-clinical context. Therefore, the question of prevalence remained open: is the ability to adopt a SCV phenotype mostly restricted to clinical isolates or clinical context, from chronic infections -or not?

Here, we investigated the distribution of a SCV-based adaptative strategy in *P. aeruginosa* by screening 22 strains from diverse origins. Selective conditions were achieved by static cultivation, a culture condition that generates different microenvironments, as seen by the formation of a pellicle biofilm at the air-liquid interface. Plating of bacteria from static cultures of all 22 strains resulted in the formation of small colonies with sizes similar to SCVs described in other studies (16, 18). However, SCVs are not exclusively defined by the smaller size of their colonies. SCVs are also often identified based on the rugosity of the colony formed on Congo Red agar plates, hence the alternate name RSCVs for <u>R</u>ugose <u>S</u>mall <u>C</u>olony <u>V</u>ariants (14, 24, 43). Nevertheless, rugosity is a subjective feature, and its description may vary according to the observer and culture conditions. Indeed, we have observed that the rugosity level varies between strains. This might be especially true for strains originating from diverse environments, as in the present study. Thus, we decided to take advantage of the various additional phenotypes described for SCVs to ascertain their identity (Fig. 6). To this end, we focused on five strains representing diverse environmental origins. Based on their phenotypic features, the small colonies obtained from each parental strain were clustered into two distinct groups. Small colonies classified in Cluster 1 shared several inter-strain phenotypic features, including reversion visible after 48h. Based on known features, these small colonies can be defined as typical SCVs, validating that SCVs emerge from *P. aeruginosa* isolated from any origins. Thus, the ability to switch to the SCV phenotype appears an intrinsic feature of the species.

**Fig. 6.**
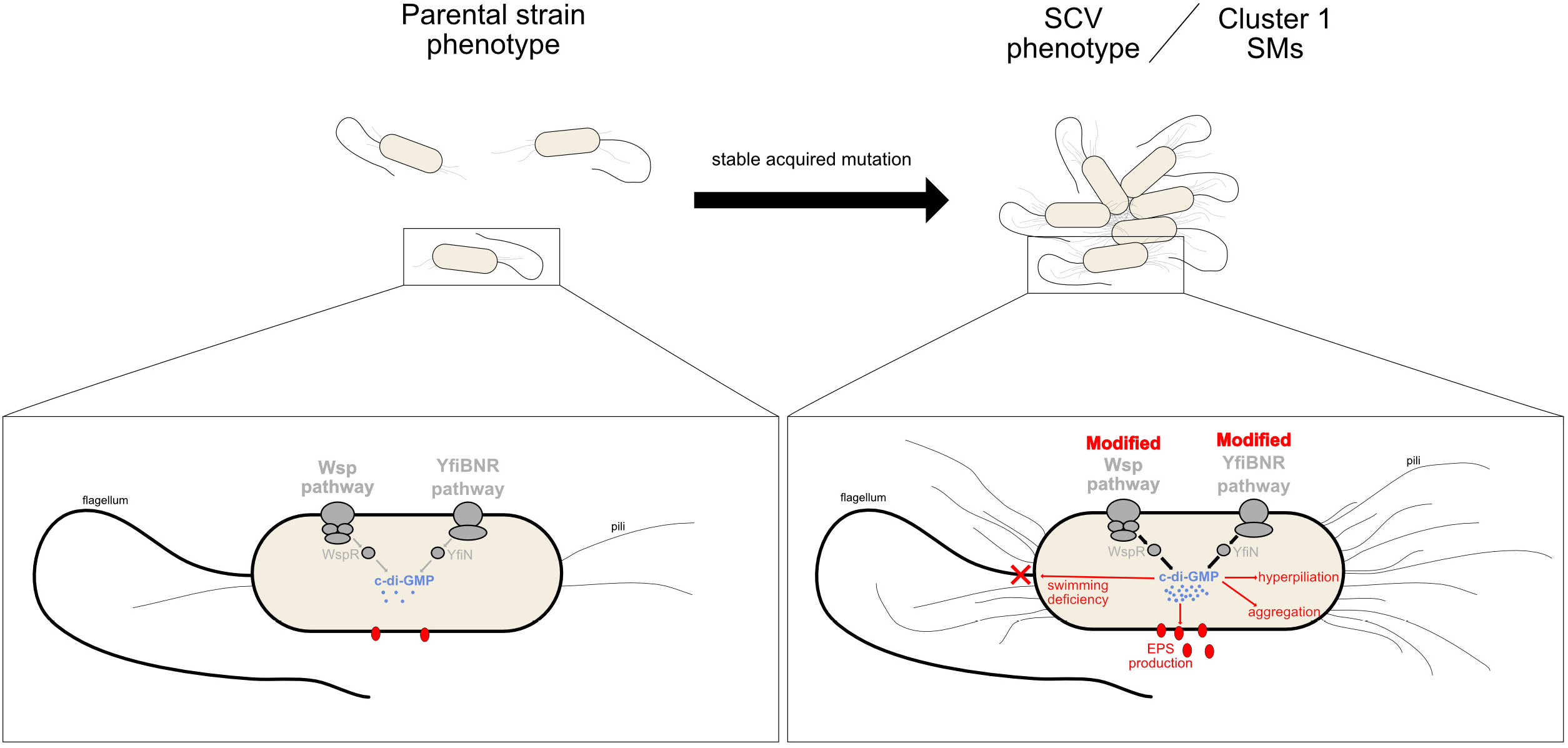
Schematic summary of SCV features compare to parental strain.

SCVs have always been isolated from biofilm-promoting conditions or from environments where biofilms thrive (16, 39, 40). SCVs are especially prone to adherence and biofilm formation (18, 36, 39). The attached mode of growth (biofilm) is a widespread lifestyle in all types of environments (46-48). Biofilms are protective barriers for their bacterial components and increase tolerance to antimicrobials as compared to free-living bacterial cells, and enhance the ability to survive in extreme conditions, such as desiccation (49-51). Thus, one can easily conceive that the switch to the SCV phenotype confers a significant advantage for colonization of various ecological niches, accounting for the apparently conserved rapid switching to the SCV phenotype.

### The SCV phenotype results from a rapidly acquired stable genetic mutation in c-di-GMP systems

Intracellular c-di-GMP levels regulate all phenotypes associated with SCVs: EPS production, motility, adherence, etc. (33-35). It is obvious that c-di-GMP thus plays a major role in the regulation of the SCV phenotype, but the role of c-di-GMP remains elusive in the mechanism of emergence. Several studies have reported that SCVs display mutations in genes involved in regulation of c-di-GMP level, particularly genes included in the YfiBNR and Wsp pathways (14, 24, 43). These mutations lead to the activation of the DGC and subsequent increase in c-di-GMP levels, either due to inactivation of the DGC repressor or constitutive activity of the chemosensory protein at the top of the signal transduction pathway. However, none of these studies were based on environmental strains, which is relevant since *P. aeruginosa* is naturally a saprophyte. Thus, involvement of c-di-GMP in the regulation and/or emergence of the SCV phenotype from an environmental parental strain remains elusive. By sequencing the whole genome of cluster 1 SCVs obtained from the prototypical clinical strain *P. aeruginosa* PA14 and the environmental strain *P. aeruginosa* 57RP, we wanted to determine if environmental strains can also switch to the SCV phenotype by using the same mechanism as PA14. We detected stable acquired mutations in all the PA14 SCVs genomes we sequenced (14, 24). However, the previous reports have concluded on the stability of the mutation in PA14 SCV by using genetic complementation and observing reversion to parental-like morphotype (14, 24). Here, we present the first study that conclude on the stability of mutations based on the whole genome sequencing of revertants, highlighting a second spontaneous mutation responsible for the switch of morphotype observed.

In a previous report, studying genetic evolution of PA14 upon infection in an animal model, Gloag *et al*. found that all PA14 SCVs displayed driver mutations in the Wsp pathway, mainly in *wspA* and few in *wspF*. Wsp is a chemosensory pathway, activated by surface sensing from WspA, and ultimately leading to the activation of the diguanylate cyclase WspR which catalyses the synthesis of c-di-GMP. This activation loop is regulated by the repressor WspF that acts to reset the system upon phosphorylation by WspE (28, 52). Here, PA14 SCVs from our standing culture conditions acquired mutations in genes of the Wsp pathway, *wspA* and *wspF*, but also in *yfiN*. The distinct membrane-integral DGC YfiN belongs to the YfiBNR pathway. YfiN activity is inversely controlled by the small periplasmic protein YfiR (repression) and the outer membrane protein YfiB (43, 53). While a previous study has reported the importance of the *yfiBNR* operon in emergence of SCVs in *P. aeruginosa* PAO1, our results confirm the importance of *yfiBNR* for SCV emergence in PA14 as well (43). The interest is emphasized by the phylogenetic distance between PAO1 and PA14 strains, belonging to two distinct phylogenetic groups of *P. aeruginosa* (54). Importantly, unlike in this previous study, the *yfiBNR* mutation in PA14 genome leading to SCV phenotype arose spontaneously. In PAO1, targeted mutation of the gene encoding the DGC inhibitor YfiR led to SCV phenotype (43). Here we present the first report of a spontaneous mutation directly in the gene coding the DGC itself, *yfiN*, leading to the SCV phenotype. Surprisingly, this mutation results in an increased c-di-GMP level, maybe due to a distinct steric hindrance following the replacement of a cysteine residue by a serine.

Apart from *P. aeruginosa* PAO1 and PA14, sometimes considered as laboratory strains rather than clinical strains, this is the first report studying the mechanism of SCV phenotype emergence in *P. aeruginosa* environmental strains, such as 57RP. By sequencing the genomes of several independently evolved 57RP SCVs, we found that the same mutation frequently occurred in *wspA*. Interestingly, the exact same *wspA* Δ285-298 deletion was also the most common mutation detected in PA14 SCVs upon infection of a murine chronic model (24). Several studies have reported mutations in this particular region (14, 55, 56), probably because this region may be hypermutable (24). Yet, it is striking that the deletion was the same in SCVs from PA14 and 57RP (24), especially since SCV selection conditions were completely very different. Since this deleted sequence is flanked by repeated inverted sequences, it could be a mobile element (24). However, we were not able to find this sequence at any other location in the parental strain genomes, nor in revertants genomes, suggesting that this may not be a reversible deletion. More so, the deletion was stable since reversion was due to a second mutation in a downstream gene of the *wsp* operon. This deletion was proposed to lead to a constitutive signaling and autoinduction of the Wsp pathway by alteration of methylation/demethylation of WspA, which would result in an increase in c-di-GMP production (24). While this is the first report of a Wsp mutation leading to SCV emergence in an environmental *P. aeruginosa* strain, it should be emphasized that a mutation in the Wsp pathway leading to the SCV phenotype was first detected in the environmental strain *P. fluorescens* Pf0-1 (55, 57). All together, these results indicate that c-di-GMP plays a central role in SCVs emergence, in strains of both clinical and environmental origins.

Interestingly, only one mutation was identified in the genomes of *P. aeruginosa* PA14 and 57RP SCVs. The sole other study which has also detected mutations having appeared spontaneously in *P. aeruginosa* PA14 SCV reported secondary mutations in the SCVs genome (24). Also, after a unique 65h incubation of the parental strain under static culture conditions, SCVs from both *P. aeruginosa* PA14 and 57RP represented 44.4 % and 18.7 % of the total population, respectively. This indicate that, regardless of the strain origin, mutants in c-di-GMP pathways are selected *in vitro* to adapt to specific conditions and switch to the SCV phenotype (Fig. 6). Among all the c-di-GMP pathways known in *P. aeruginosa*, Wsp and YfiBNR seem to be preferred pathways involved in SCV emergence. However, although the c-di-GMP increase resulting from alteration in Wsp and YfiBNR pathways are responsible for SCVs emergence, the opposite was not true. Inactivation of the DGCs WspR and YfiN, resulting in inability to produce c-di-GMP through this pathway did not affect the rate of emergence of SCVs. Thus, regulation of SCV emergence through c-di-GMP mechanisms could be based on interchangeable DGCs, ready to take over the inactivity of one of the pathways.

### SCVs could also emerge from a phase variation mechanism, undetectable under laboratory conditions

Phase variation is a common phenomenon among Gram-negative bacteria and is typical of bacteria thriving in heterogeneous ecological niches (21, 22, 58), notably *P. aeruginosa* (19). Unlike acquisition of stable mutations, phase variation mechanism represents a significant advantage for the rapid adaptation to sudden changes in the environment (59, 60). Indeed, phase variation mechanisms lead to emergence of a heterogeneous population in which the best suitable phenotype will multiply until the conditions fluctuate again and the selected phenotypes revert to another phenotype.

Although SCVs are due to stable genetic mutations under our experimental conditions, i.e. irreversible mutations, we cannot exclude that the adoption of the SCV phenotype could also rise from a reversible phase variation regulated mechanism in natural habitats. Several reports support this hypothesis. First, phenotypes traditionally related to SCV (motility, aggregation) are often regulated by phase variation mechanisms (21). In addition, reversible adaptation mechanisms are based on transitory DNA rearrangements (gene conversion, genomic inversion, DNA recombination..) and lead to variation in gene expression (20). Indeed, one recent study reports a large genomic inversion in *P. aeruginosa* SCVs (61). Finally, reversion of SCVs has been observed several times and could be due to phase variation instead of emergence of a second mutation, but no whole genome sequencing has been performed to conclude (15, 16, 18). Yet, SCVs reversion occurred toward a phenotype likely different from the parental morphotype (16, 23), suggesting that regulation is not necessarily an ON/OFF switch on a particular locus and could be due to a secondary mutation in the genome.

Under our conditions, SCVs from phase variation mechanism could have arisen and were undetectable in our conditions with our technique. Two elements could explain this limitation: (1) the “reversible” SCVs are present but in undetectable quantities to be observed after sampling and agar spreading or (2) “reversible” SCVs were present but reverted to another morphotype when the samples were spread on agar plates. Indeed, various phenotypes were observed on agar plates after 65h of standing incubation, likely to have emerged in the static liquid culture but they could have also emerged directly on the agar plate. To verify this hypothesis and verify that SCVs can also arise from a reversible mechanism, it would be interesting to follow the emergence of SCVs in the static culture using a detectable marker.

### Small colonies are not necessarily SCVs, nor variants

During our experiments with static cultures, we observed several small-colony morphotypes. Based on our PCoA analysis, a proportion of them were clustered in two distinct groups (Fig. 2). Except for strain 57RP, the SMs from Cluster 2 did not display clear reversion after 48 h on solid medium (data not shown). However, SMs from Cluster 2 could still be able to revert in conditions outside the ones tested in our study. Also, their rate of emergence seemed too high for mutants (Table 1). Thus, we wonder if cluster 2 SMs should be identified as variants based on our criteria.

In contrast with SMs from Cluster 1, SMs from Cluster 2 showed inter-strain heterogeneous features. We observed a large diversity of morphotypes on plates prepared from our static cultures. Among them, large colonies also displayed features similar to revertants (16). This observation indicates that reversion could have occurred in the static liquid cultures, and intermediate forms could consequently be isolated. Maybe several mechanisms are acting in parallel to induce the phenotypical diversity we observed, thus promoting the selection of the best adapted subpopulation.

The SCV phenotype has been linked to the persistence of *P. aeruginosa* in the context of infections in a human host, notably because of its increased resistance against antimicrobials and host immunity. However, we have demonstrated here that strains isolated from soil, food and hospital environments can also readily adopt a SCV phenotype. This indicates that the ability of *P. aeruginosa* to form SCVs is a conserved feature of this species, and SCVs emergence is not exclusively related to the pressure of the infection-related clinical environment. This is the first report of high prevalence of SCVs among *P. aeruginosa* strains, regardless of the origin of the isolates. The SCVs identified showed specific mutations in genes related to regulation of the c-di-GMP intracellular levels. The Wsp and YfiBNR systems were the primary pathways used to increase c-di-GMP level and switch to SCV phenotype. Emergence of SCV in the various habitats allows *P. aeruginosa* to rapidly adapt and persist under diverse environmental conditions, accounting for its versatility and persistence. A deeper comprehension of the adaptation strategy used by *P. aeruginosa* could ultimately provide innovative strategies for eradication of this opportunistic pathogen of public concern.

## MATERIALS AND METHODS

### Bacterial strains and growth conditions

Bacterial strains are listed in Table 1 and their specific origin are listed in Table S2. In this study, the term “parental strain” designs the original strain used to evolve other morphotypes in static cultures, including SCVs. Strains were grown in tryptic soy broth (TSB; BD), at 37°C in a TC-7 roller drum (New Brunswick Scientific) at 240 rpm for the parental strains and at 30°C in an Infors incubator (Multitron Pro) at 180 rpm (angled tubes) for the isolated evolved morphotypes. Static cultures were inoculated with the parental strain at an initial OD_600_ of 0.05 and incubated at 30, 30.9, 32.2, 33.9, 36.3, 38, or 40°C for 65 hours. Cultures were then spread on tryptic soy 2% agar plates (TS-Agar; AlphaBiosciences), unless stated otherwise. Two percent agar was utilized to limit expansion of colonies and improve isolation of the distinct morphotypes.

### Bradford protein assay

Due to the highly aggregative properties of SCVs, OD_600_ measurements were not appropriate to evaluate growth of some of the isolated evolved morphotypes. Instead, the Bradford protein assay was used to quantify the concentration of total proteins in all our samples. Pellets from 1 ml of culture were resuspended in 1 ml 0.1 N NaOH and incubated 1 h at 70°C. Protein concentrations were measured on samples according to the manufacturer guidelines for the Bradford reagent (Alfa Aesar).

### Phenotypic tests

Overnight (O/N) cultures of parental strains and their isolated morphotypes were grown at 30°C in an Infors incubator (Multitron Pro) at 180 rpm in angled tubes. Since biofilm formation occurred in cultures, they were transferred to clean tubes to perform experiments or Bradford protein quantifications. Statistical analyses were achieved using Ordinary one-way analysis of variance (ANOVA). Each phenotypic test was performed in technical triplicates.

### Morphology on Congo red plates

A 1% Congo red solution in water (Fisher SCIENTIFIC) was added to TS-Agar 2% to a final concentration of 0.1%. Ten µL of culture were spotted on the plates. Plates were incubated at 30°C and observed after 24 h, 48 h and 96 h. Plates were observed with a binocular StemiDV4 (Zeiss) and photos were taken with the camera DMC-ZS60 (Panasomic Lumix).

### Swimming motility tests

Swim plates (20 mM NH_4_Cl, 12 mM Na_2_HPO_4_, 22 mM KH_2_PO_4_, 8.6 mM NaCl, 0.5% Casamino acids (CAA), 0.3% Bacto-Agar (BD), supplemented with 1 mM MgSO_4_, 1 mM CaCl_2_ and 11 mM dextrose) were prepared and dried for 15 min under the flow of a Biosafety Cabinet. A volume of 2.5 µL of culture was inoculated in the agar. Plates were incubated for 20 hours at

30°C. Swimming ability was assessed by measuring the area (mm^2^) of the turbid circular zone using ImageJ. All experiments were performed in triplicates.

### Biofilm formation

Microtiter (96-well) plates containing 1/10 TSB supplemented with 0.5% CAA were inoculated from a transferred O/N culture in order to obtain a starting concentration of 70 mM proteins. Each sample was inoculated in five different wells. Plates were incubated at 30°C without agitation. After 6 and 24 h, plates were rinsed thoroughly with distilled water and 200 µL of a 1% crystal violet solution was added to each well. After 15 minutes of incubation at room temperature, plates were rinsed thoroughly with distilled water and the dye was solubilized in 300 µL in 30% acetic acid. The absorbance was measured at 595 nm with a microplate reader (Cytation3, Biotek). Initiation of biofilm formation was calculated as the % of biofilm formed after 6 h of incubation compared with total biofilm formed after 24 h incubation. Total biomass of the biofilm was calculated as the amount of biofilm formed after 24 h, measured by crystal violet absorbance at 595 nm after 24 h of incubation.

### Pyoverdine production

Overproduction of pyoverdine was previously noted as a feature of strain 57RP SCVs (18). We confirmed that a SCV from PA14 expresses high fluorescence level at the wavelength of pyoverdine emission, likely to account for cell aggregation and EPS overproduction. An SCV isolated from a PA14 *pvdD* mutant (62), which is no longer able to produce pyoverdine, showed lower fluorescence levels, similar to parental colonies, confirming that [1] pyoverdine production is responsible for the fluorescence detected and [2] measured fluorescence is correlated with SCV aggregation properties (Fig. S3). To measure pyoverdine production, black 96-well plates (Greiner) were filled with 200 µL of culture. Fluorescence was measured at wavelengths 390 nm/530 nm excitation/emission using a multimode microplate reader (Cytation3, Biotek).

### C-di-GMP quantification

Intracellular levels of c-di-GMP were assessed with the fluorescence-based biosensor pCdrA-gfpC (63, 64), acquired as addgene plasmid #111614; http://n2t.net/addgene:111614 ; RRID:Addgene_111614. Purified plasmids were transformed by electroporation in evolved morphotypes obtained from static cultures (65). Transformants were selected on TS-Agar 2% supplemented with 100 µg/ml gentamycin. Three clones for each transformed morphotypes were cultured in TSB supplemented with gentamycin 100 µg/ml. Cultures were washed twice in fresh TSB to get rid of a potential non-specific fluorescence due to secreted fluorescent pigments as pyoverdine. Fluorescence was measured using a Cytation3 microplate reader (BioTek) at 490 nm/515 nm (excitation/emission) in black 96-well plates (Greiner). The non-transformed strain was used as a control. Fluorescence from the control was subtracted from the fluorescence signal for the transformed strains.

### PCoA analysis

Colonies identified as SMs compared with their parental isolate (see Results) were used to perform a principal coordinate analysis (PCoA). Statistical analyses were performed using RStudio software version 1.3.1093 (66) with normalised data showed in Table S1. A Euclidean distance matrix was used to generate a clustering of the bacterial isolates according to their phenotypical profile. A Similarity Profile Analysis (simprof) was performed to determine the number of significant clusters produced using hclust with the assumption of no *a priori* groups. Significant clusters were considerate when at least two evolved morphotypes constituted it.

### Sequencing and analysis

To analyse SCVs genomes and compare them with corresponding parental strain genomes, genomic DNA was extracted using the EasyPure® Genomic DNA Kit (Transgen Biotech), according to the manufacturer’s protocol from 200 µL of O/N culture of colony variants in TSB. Clonal DNA was sequenced on the Illumina NextSeq 2000 at the Microbial Genome Sequencing center (MiGS); 2×151 bp sequencing paired end reads were trimmed and quality filtered using fastp v0.23.2 (67). Filtered reads were assembled using Skesa through Shovill v1.1.0 and annotated using prokka v1.14.6 to be used as reference sequences (68). The reads were then used for variant calling using snippy v4.6.0 (https://github.com/tseemann/snippy) using default settings.

Presence of a 42 bp deletion in *wspA* for additional 57RP colony variants DNA was confirmed by amplifying a 200 bp PCR fragment with primers F-CGGAGACTTCGCTCATGGT and R-AGAGCTCAAGGGCCTGGT. The detection of amplified products was performed on a 2% agarose gel electrophoresis.

## ACKNOWLEDGMENTS

We thank Cynthia Bérubé for her help with the c-di-GMP biosensor preliminary experiments, and Thays de Oliveira Pereira for critical reading of the manuscript.

This work was supported by grant MOP-142466 from the Canadian Institutes of Health Research (CIHR). Dr. Alison Besse was a Fellow of the post-doctoral grant Calmette and Yersin from the Institut Pasteur.

The funders had no role in study design, data collection and interpretation, or the decision to submit the work for publication.

AB and ED conceived the project, contributed to experimental design and interpreted results. AB, MCG and MT contributed to data acquisition. AV and AB analysed sequencing. AB wrote the manuscript. MCG and ED reviewed and edited the manuscript.

